## Supplementary File for "Predicting bacterial phenotypic traits through improved machine learning using high-quality, curated datasets"

Julia Koblitz *et al.*

**This PDF file includes:**

Figs. S1 to S4
Tables S1 to S1

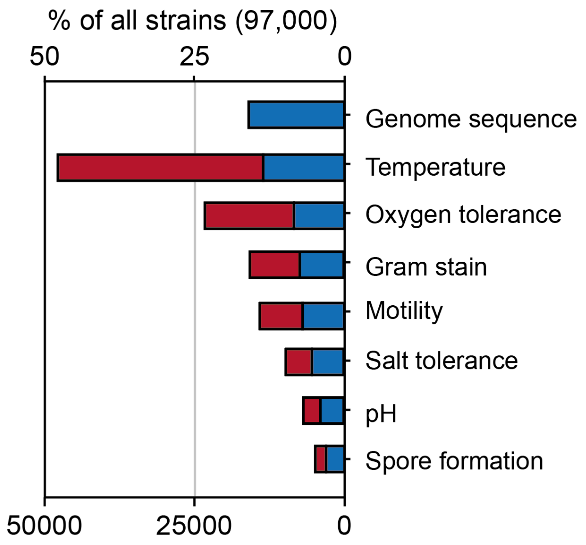


**Figure S1: Availability of genome sequences and most common types of phenotypic information for all bacterial strains covered by the Bac*Dive* database**. In contrast to figure 1 of the main manuscript, the strains are not limited to type strains. Records are colored blue if a genome is present and red if not.

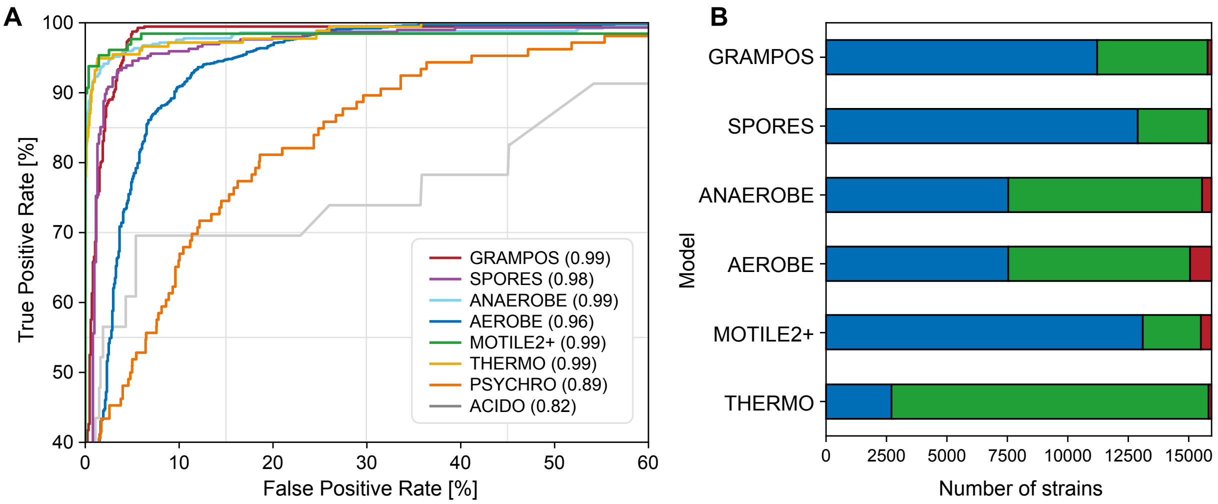


**Figure S2: Model performance metrics and improvement of the Bac*Dive* database with the models from this study. (A)** ROC curves indicate model performance. The legend also contains the calculated AUC. Axes are limited to minimum 40% true positive rate and maximum 60% false positive rate. **(B)** The number of strains to which prediction data has been added. Blue: new data where the strain did not have information on the trait before, green: consistent data where the predicted label is consistent with the manually curated information, red: contradicting data where the prediction contradicts the curated data.

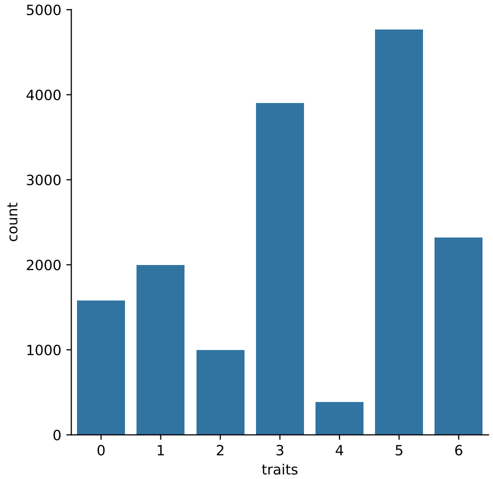


**Figure S3: Number of new traits that were added to strains in Bac*Dive*.** 0 means no new datasets were added to a strain, 7 means the strain had none of the phenotypic data before.

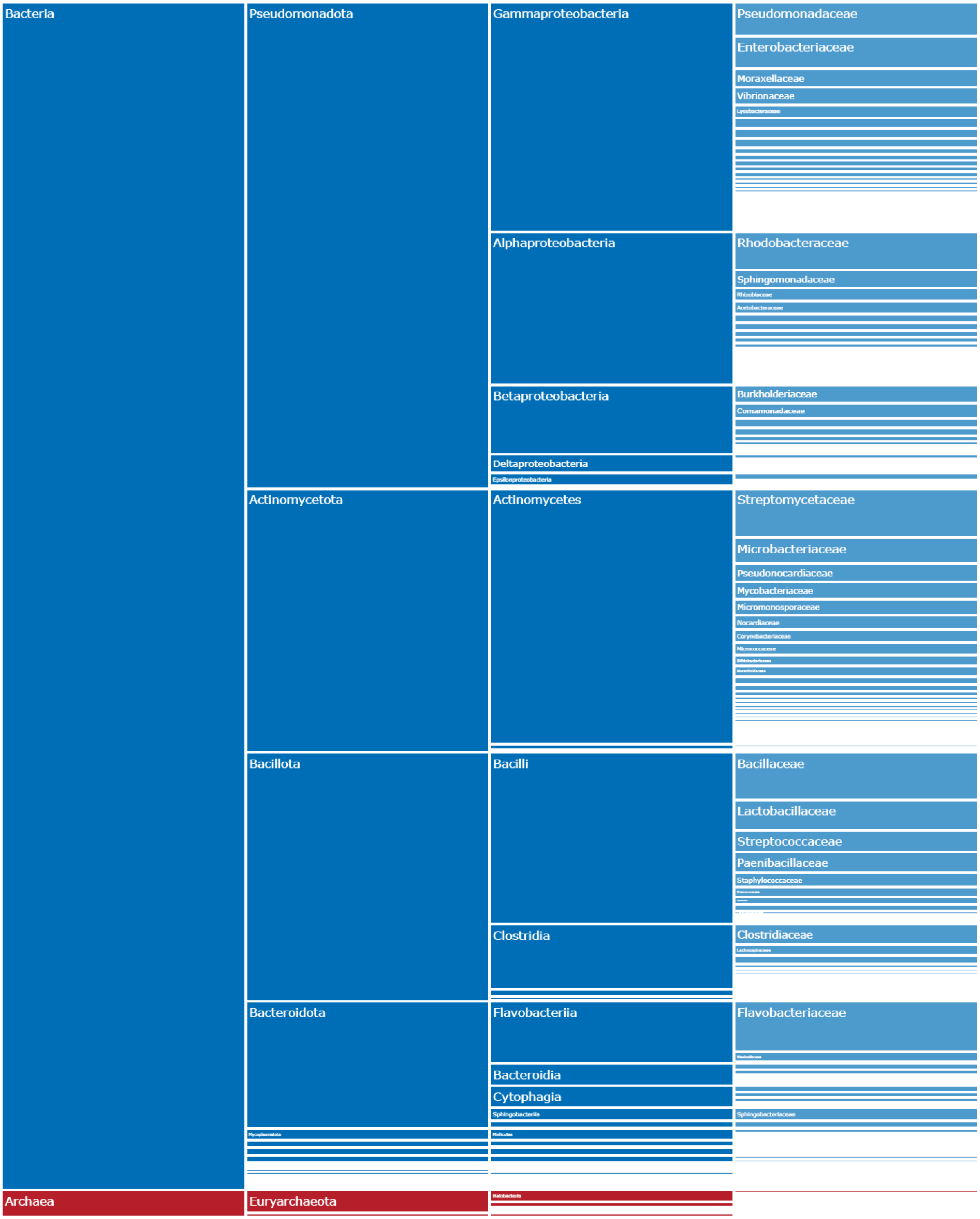


**Figure S4: Taxonomic classification of all strains used in the training and test dataset.** Taxonomic names according to LPSN.

**Table S1:** Feature importances of the MOTILE model.

| Pfam | Description | Importance |
|---|---|---|
| PF00771 | FHIPEP family | 0.024 |
| PF02154 | Flagellar motor switch protein FliM | 0.023 |
| PF06429 | Flagellar basal body rod FlgEFG protein C-terminal | 0.022 |
| PF01514 | Secretory protein of YscJ/FliF family | 0.021 |
| PF00669 | Bacterial flagellin N-terminal helical region | 0.020 |
| PF14841 | FliG middle domain | 0.020 |
| PF02049 | Flagellar hook-basal body complex protein FliE | 0.019 |
| PF01311 | Bacterial export proteins, family 1 | 0.018 |
| PF00813 | FliP family | 0.015 |
| PF01313 | Bacterial export proteins, family 3 | 0.014 |
| PF07195 | Flagellar hook-associated protein 2 C-terminus | 0.013 |
| PF01052 | Type III flagellar switch regulator (C-ring) FliN C-term | 0.013 |
| PF00015 | Methyl-accepting chemotaxis protein (MCP) signalling domain | 0.013 |
| PF14842 | FliG N-terminal domain | 0.013 |
| PF01312 | FlhB HrpN YscU SpaS Family | 0.012 |
| PF01706 | FliG C-terminal domain | 0.012 |
| PF01584 | CheW-like domain | 0.011 |
| PF00700 | Bacterial flagellin C-terminal helical region | 0.011 |
| PF18269 | T3SS EscN ATPase C-terminal domain | 0.009 |
| PF08345 | Flagellar M-ring protein C-terminal | 0.009 |
| PF03963 | Flagellar hook capping protein - N-terminal region | 0.008 |
| PF00033 | Cytochrome b/b6/petB | 0.008 |
| PF02119 | Flagellar P-ring protein | 0.007 |
| PF06798 | PrkA serine protein kinase C-terminal domain | 0.007 |
| PF02561 | Flagellar protein FliS | 0.007 |

**Table S2:** Mobility test of selected strains with 21 flagellum Pfams that were reported as non-motile in the literature.

| Strain | Bac*Dive*-ID | Confidence (MOTILE_2) | Motility | Reference |
|---|---|---|---|---|
| *Heyndrickxia oleronia* DSM 9356 | 1102 | 84.6 | ++ | (*32*) |
| *Halobacillus kuroshimensis* DSM 18393 | 1349 | 84.7 | + | (*33*) |
| *Salipaludibacillus neizhouensis* DSM 19794 | 1293 | 83.0 | (+) | (*34*, *35*) |
| *Terribacillus halophilus* DSM 21620 | 1518 | 81.9 | (+) | (*36*) |
| *Priestia endophytica* DSM 13796 | 1203 | 78.6 | - | (*37*) |
